# The GRASP domain in Golgi Reassembly and Stacking Proteins: differences and similarities between lower and higher Eukaryotes

**DOI:** 10.1101/522573

**Authors:** Luís F. S. Mendes, Natália A. Fontana, Carolina G. Oliveira, Marjorie C. L. C Freire, José L. S. Lopes, Fernando A. Melo, Antonio J. Costa-Filho

## Abstract

The Golgi complex is part of the endomembrane system and is responsible for receiving transport cargos from the endoplasmic reticulum and for sorting and targeting them to their final destination. To perform its function in higher eukaryotic cells, the Golgi needs to be correctly assembled as a flatted membrane sandwich kept together by a protein matrix. The correct mechanism controlling the Golgi cisternae assembly is not yet known, but it is already accepted that the Golgi Reassembly and Stacking Protein (GRASP) is a main component of the Golgi protein matrix. Unlike mammalian cells, which have two GRASP genes, lower eukaryotes present only one gene and distinct Golgi cisternae assembly. In this study, we performed a set of biophysical studies to get insights on both human GRASP55 and GRASP65 and compare them with GRASPs from lower eukaryotes (*S. cerevisiae* and *C. neoformans*). Our data suggest that both human GRASPs are essentially different from each other and GRASP65 is more similar to the subgroup of GRASPs from lower eukaryotes. GRASP55 is present mainly in the Golgi medial and trans faces, which are absent in both funguses, while GRASP65 is located in the cis-Golgi. We suggest that the GRASP65 gene is more ancient and the paralogue GRASP55 might have appeared latter in evolution, together with the medial and trans Golgi faces in mammalians.

## 1. INTRODUCTION

The Golgi complex is part of the endomembrane system and is responsible for receiving transport cargos from the endoplasmic reticulum to sort and to target them to their final destination [1,2]. Furthermore, this organelle works as the main glycosylation hub inside the cell and is also responsible for many lipid syntheses [1,2]. In order to achieve such an expressive number of functionalities in higher eukaryotic cells, the Golgi needs to be correctly assembled as a flattened membrane sandwich kept together by a protein matrix [1]. The disruption of this assembly ultimately leads to failure in correct protein glycosylation and sorting, besides protein secretion impairment [1,3,4,5].

The exact mechanism controlling the Golgi cisternae assembly is currently unknown, but it has been largely accepted the pivotal role of the Golgi Reassembly and Stacking Protein (GRASP) as main part of the Golgi protein matrix [6,7,8]. There are two paralogue genes found in mammalians (GRASP55 and GRASP65) which work in synergy to keep both cisternae stacking and their lateral tethering, thus building the Golgi ribbon [8,9]. It has been previously observed that the double knockout of GRASP55 and GRASP65 by CRISPR-Cas9 technology led to a total Golgi fragmentation [10]. More than that, GRASP silencing also led to incorrect protein glycosylation [3] and accelerated protein transport to the plasma membrane [4]. Both GRASP proteins require a double membrane anchoring to promote their correct trans dimerization, responsible for bringing the Golgi cisternae together [11]. While GRASP65 is located mainly in the cis part of the Golgi, GRASP55 is present in the medial and trans faces [6,7]. These unique localizations of both GRASPs have been explored as Golgi markers for co-localization experiments in several manuscripts [12,13].

GRASPs are observed in all eukaryotic cells with the exception of plants, where no obvious homologues have been found so far [9,14]. Interestingly, mammalian cells have two GRASP genes [14], while in lower eukaryotic cells, where the correct Golgi cisternae assembly is not a common feature, only one gene has been reported [14]. *Saccharomyces cerevisiae* has one GRASP gene and only 40% of its cisternae are assembled into stacks, whereas in the human pathogen *Cryptococcus neoformans*, which also has one GRASP gene, no Golgi stacks are observed [15,16]. Curiously, *Plasmodium falciparum*, the causative agent of malaria, has in its DNA one GRASP gene that encodes for two GRASP isoforms coming from a splice variant [17]. In this system, the first GRASP relies on a well-conserved myristoylation motif typical of the human GRASPs, while the variant displays a different N-terminus, similar to GRASPs found in fungi [17].

Even tough GRASPs were initially found as structural factors of the Golgi complex [14], an unusual (and still unclear) participation in unconventional protein secretion (UPS) of several proteins has been observed [18,19]. GRASPs have a fundamental involvement in the stress-induced UPS (more specifically, UPS types 3 and 4 [19]) but their exact role is still a matter of intense debate [19]. A recent paper has suggested that GRASPs act as a tethering factor of autophagosome accessory proteins needed for the autophagosome-lysosome fusion and also on autophagosome maturation under glucose deprivation [20,21]. Since nutrient starvation is one of the main activators of type 3 UPS, it has been suggested that GRASP might be recruited by the autophagosome and by the plasma membrane to enable the release of the UPS cargo, but this hypothesis is still to be proven. A previous work has suggested a direct (and necessary) interaction of GRASPs with the protein to be secreted, especially in the case of type 4 UPS [22]. This direct interaction with the protein cargo has already been observed in other situations [23,24,25,26]. Some specific roles have also emerged, such as the participation in the formation of specific autophagosome-like structures (named compartments of unconventional protein secretion, or CUPS), but regardless their true nature and existence in yeast [27,28], so far there are no evidences that these structures exist in mammalian cells [29].

The GRASP structure is composed of two main regions, a conserved N-terminal portion with two protozoa-like PDZ domains, called GRASP domain, and a non-conserved C-terminus, rich in serine and proline residues and presenting an intrinsically disordered behavior [14,30]. The GRASP domain is the “functional” region of this protein, acting in the dimerization step during cisternae stacking and also “grasping” protein partners in an array of different situations [9,14,31]. The C-terminus, also known as SPR domain, has a regulatory functionality by means of its phosphorylation [14], cleavage sites [32] and O-GlcNAcylation in nutrient-rich conditions [33]. It has been previously reported that the full-length GRASP from *Cryptococcus neoformans* behaves as a molten globule-like protein, with an overall structural dynamic in μs-ms timescale and high structural plasticity [30]. The physicochemical perturbations in the surroundings of the biological membrane seem to be the main inducer of disorder-to-order transitions in GRASP [34].

Despite the great number of studies focusing on GRASPs in both biological and biophysical context, several aspects remain elusive. For instance, although GRASPs are conserved among the GRASP-containing organisms, there are no obvious reason of why there are two GRASPs in mammalians and only one in lower eukaryotes, and whether GRASP55 and GRASP65 show the same behavior in solution although having distinct functionalities and protein partners [9,14]. Whether or not the variations seen in the number and behavior of GRASP homologues correlates with the significant variability in the Golgi structure or with the particularities in UPS among different organisms still deserves more attention. In this study, we performed a set of biophysical studies on the GRASP domains of both human GRASP55 and GRASP65 (named DGRASP55 and DGRASP65, respectively) and compare them with two members of GRASPs from lower eukaryotes (*S. cerevisiae* and *C. neoformans*) to get insights on the structural behavior of GRASPs from different system complexity.

## 2. MATERIALS AND METHODS

### 2.1. Protein expression and purification

*C. neoformans* GRASP domain (CnDGRASP) and *S. cerevisiae* GRASP domain (ScDGRASP) were expressed and purified as discussed elsewhere [30,35]. Human GRASP55 and GRASP65 genes were synthesized with a codon-optimization for *E. coli* expression and the GRASP domain of GRASP55 (1-208) and of GRASP65 (1-211) were amplified by PCR and subcloned into the pWALDO-d vector using the Nde1 and BamH1 sites and further transformed into Rosetta (DE3) cells. Liquid inoculum of LB-medium was used for cell growth at 37° c until OD_600_ nm of 1.0. Protein expression was induced by adding Isopropyl-β-D-thiogalactoside at a final concentration of 0.5 mM for 18 hours at 20 c and 220 rpm. Cells were sedimented by centrifugation (7.000xg per 5 minutes at 4°c) and resuspended in a solution of 20 mM Tris-HCl, 150 mM NaCl, 10 mM 2-Mercaptoethanol, 0.5% Triton X-100 and 250 μM PMSF, pH 8.0. Cells were disrupted by sonication and clarified by centrifugation (18.000xg per 20 minutes at 4°c). The soluble fraction was loaded into a Ni-NTA column, and the bound protein was washed with 20 mM Tris-HCl, 150 mM NaCl, 10 mM 2-Mercaptoethanol, pH 8.0 (buffer A), buffer A + 20 mM imidazole, buffer A + 50 mM imidazole and eluted with buffer A + 500 mM imidazole. Two miligrams of purified TEV were added for each litter of expressed GRASP55 or GRASP65 GRASP domain (DGRASP55 and DGRASP65, respectively, from now on) and the final solution (around 15 mL) was dialyzed against 4 L of 20 mM Tris-HCl pH 8.0, 50 mM NaCl and 7 mM 2-Mercaptoethanol, overnight at 4°c. The solution was then reloaded into a Ni-NTA column and the super flow was collected, concentrated and applied in a Mono Q 5/50 GL (GE Healthcare Life Sciences) coupled to an *Äkta purifier* system (GE Healthcare). The protein solution was submitted to a linear gradient of NaCl from 50 to 350 mM (25 mM Tris-HCl pH 8.0, 10 mM 2-Mercaptoethanol). The fractions corresponding to pure DGRASP55/65 were then concentrated and applied into a Superdex200 10/300 GL gel filtration column (GE Healthcare Life Sciences) equilibrated in a 25 mM Tris-HCl pH 8.0, 150 mM NaCl, 7 mM 2-Mercaptoethanol. For protein concentration, an Amicon Ultra-15 Centrifugal Filter with a NMWL of 10 kDa (Merck Millipore) was used. Protein concentration was measured using the extinction coefficient at 280 nm calculated using protparam webserver [36] considering the non-native aminoacids present at the C-terminus after TEV cleavage. For DGRASP55 the *ε*_280 *nm*_ = 28,420 M^−1^cm^−1^ and Abs 0.1% (=1 mg/ml) 1.199, while for DGRASP65 is *ε*_280 *nm*_ = 28,420 M^−1^cm^−1^; Abs 0.1% (=1 mg/ml) 1.182 were used. For CnDGRASP and ScDGRASP the parameters were as presented elsewhere [30,35].

### 2.2. Circular Dichroism (CD)

Conventional far-UV CD experiments were performed in a Jasco J-815 CD Spectrometer (JASCO Corporation, Japan) equipped with a Peltier temperature control and using a quartz cell with a 1 mm pathlength. The spectra were recorded from 260 to 198 nm, with a scanning speed of 100 nm·min^−1^, spectral bandwidth of 1 nm, response time of 0.5 s and taking individual measurements prior to the average in order to check protein instability after the final spectra. All the protein stock solutions were at a minimum concentration where the dilution in a 20 mM sodium phosphate pH 8.0 (or any desired buffer) was at least 20-fold. Urea denaturation was carried out in a 0.8 M urea steps with an overnight incubation at 4°C and a one-hour incubation at 25°C prior to the measurements. Experiments with pH variation were performed with the same criteria of dilution in a 20 mM glycine-HCl (pH 3.0), 20 mM sodium acetate/acetic acid (pH 4.0), 20 mM sodium phosphate (pH 5.0-8.0) or 20 mM glycine-NaOH (pH 9.0 and 10.0) with an incubation time of 10 minutes at 25°C. All the experiments were performed at 25°C CD spectra reconstruction was performed using PDB2CD [37].

### 2.3. Synchrotron Radiation Circular Dichroism (SRCD) spectroscopy

SRCD spectroscopy was employed to access the lower wavelength UV region with an improved signal-to-noise ratio and to characterize conformational changes (significant or subtle) with higher accuracy than the conventional method. The SRCD spectra of the proteins in aqueous solution were collected at the AU-CD beamline at the Institute for Storage Ring facilities (ASTRID2 synchrotron), University of Aarhus, Denmark. Measurements were performed taking 3 successive scans over the wavelength range from 280 to 180 nm, using in 1 nm interval and 2 s dwell time, at 25 °C, using a 0.01027 cm pathlength cylindrical Suprasil quartz cell (Hellma Ltd.). Additionally, aliquots (0.7 nmol) of each protein were deposited on the surface of a quartz-glass plate to form a partially dried film of the proteins, promoting the evaporation of the solution under moderate vacuum. The SRCD spectrum of the dehydrated film was collected over the range 280-160 nm, with 1 nm interval and 2 s dwell time, at 25°C, taking four different rotations of the plate (0, 90, 180, and 270 degrees) in order to avoid linear dichroism effect. CDtoolX [38] software was used for all SRCD data processing, which consisted of performing the average of the three individual scans collected of each sample, baseline subtraction (spectra taken of each corresponding buffer with all additives, except for the protein), zeroing at the region at 263-270 nm, smoothing with a Savitky-Golay filter, calibration against a spectrum of a camphoursulphonic acid standard taken at the beginning of the data collection, and final data expressed in delta epsilon units. CD spectra deconvolution was performed using the webserver DICHROWEB [39,40] and selecting the reference set SP175 optimized for 190-240 nm.

### 2.4. Fluorescence Spectroscopy

Steady-state fluorescence was monitored using a Hitachi F-7000 spectrofluorometer equipped with a 150 W xenon arc lamp. The excitation and emission monochromators were set at 5 nm slit width in all tryptophan experiments and 5 nm excitation with 10 nm emission for the ANS experiments. The protein concentrations were 15 μM. For tryptophan fluorescence experiments, the selective tryptophan excitation wavelength was set at 295 nm and the emission spectra were measured from 310 up to 450 nm. For the 1-anilino-8-naphthalenesulfonic acid (ANS – 250 μM) fluorescence experiments, the excitation wavelength was set at 360 nm and the emission spectra were measured from 400 up to 650 nm. Fluorescence quenching using the water-soluble acrylamide as a quencher was performed in a serial dilution from an 8 M acrylamide stock solution with 5 minutes thermal equilibration. All the experiments were performed at 25°C

### 2.5. Bioinformatic analyses

Multiple sequence alignment using seeded guide trees and HMM profile-profile techniques were performed using Clustal Omega [41]. The evolutionary history was inferred by using the Maximum Likelihood method and JTT matrix-based model [42]. The bootstrap consensus tree inferred from 10,000 replicates is taken to represent the evolutionary history of the taxa analyzed. Branches corresponding to partitions reproduced in less than 50% bootstrap replicates are collapsed. The percentage of replicate trees in which the associated taxa clustered together in the bootstrap test (10,000 replicates) are shown next to the branches. Initial tree(s) for the heuristic search were obtained automatically by applying Neighbor-Join and BioNJ algorithms to a matrix of pairwise distances estimated using a JTT model, and then selecting the topology with superior log likelihood value. Evolutionary analyses were conducted in MEGA X [43]. Protein theoretical pI was estimated using Protparam webserver [36]. Protein structure homology-modelling was performed with Swiss-MODEL [44] and the sequences used are presented in the supplementary material.

### 2.6. Limited proteolysis

Proteolysis sensitivity was assessed using trypsin from bovine pancreas (Sigma) in a molar ratio (protein/trypsin) of 100/1 at 25°C. Ovoalbumin (GE Life Science) was used as model of well-structured proteins for direct comparison. Adding SDS-PAGE buffer and heating at 95°C for 10 minutes quenched the reactions. All reactions were checked by Coomassie stained 15% SDS-PAGE.

### 2.7. Nuclear Magnetic Resonance (NMR)

The NMR spectra of ^15^N-DGRASPs (concentration around 100 μM) were recorded at 25°C on an NMR spectrometer equipped with cryogenically cooled z-gradient probe operating at ^1^H frequency of 600 MHz (Bruker BioSpin GmbH, Rheinstetten, Germany) and using a Shigemi 5 mm symmetrical NMR microtube assembly (Sigma Aldrich). Protein expression in minimum media supplemented with ^15^NH_4_Cl was performed using a protocol presented elsewhere [45]. Cell induction and protein purification were performed using the same protocol described earlier for the unlabeled samples, except for a change in the solution used in the gel filtration step, where now a 25 mM HEPES/NaOH, 100 mM NaCl, 7 mM 2-Mercaptoethanol, pH 7.0 was used. A final volume of 5% D_2_O was added to each sample. The data were processed and analyzed using TopSpin 3.5.

## 3. Results and Discussions

### 3.1 Secondary structure comparison

The GRASP domain is composed of two PDZ domains connected in tandem [46,47]. PDZs work as protein interaction domains that bind, in a sequence-specific manner, to C-terminal peptides or internal disordered regions [48]. This domain has a very particular secondary structure organization formed by a ββ_2_βαββα_2_β_6_ arrangement, which collapses in a modular structure and where the binding grove is formed by both β_2_ and α_2_ [46,48]. However, the first GRASP domain structure determined for a member of the GRASP family (DGRASP55) showed that, unlike other eukaryotic PDZ domains, each binding groove was formed by the final (β_6_) rather than the second β-strand within the fold [46]. This is a very unusual structural organization for a PDZ domain in eukaryotes but a common fold in prokaryotes [46]. Therefore, GRASPs are formed by structurally arranged unique PDZ domains in eukaryotes, which could explain some of the unusual biophysical features observed previously [30,34,35].

The first comparison within the set of GRASP domains under investigation was based on their secondary structure organization, aiming to check for differences and/or conservation. The 3D structures of human DGRASP55 (PDB ID 3RLE), a peptide-bound human DGRASP65 and also the homologue in rat of apo DGRASP65 (PDB IDs 4KFV and 4REY, respectively) have been previously reported [46,47,49]. Here, we constructed a model of ScDGRASP and CnDGRASP using Swiss-MODEL and then aligned the four GRASP domain structures using Pymol (Figure 1A). The structures aligned very well, suggesting that they all share the same two-circular permutation PDZ fold, although the aminoacid sequence only shares around 30% identity across species (Figure 1B). The CD spectra show that they all might share a similar secondary structure content, except for CnDGRASP which presents the most pronounced spectral differences, especially in the region bellow 210 nm (Figure 1C). Because of the higher magnitude of the minimum and its shift to lower wavelengths (from 208 to 205 nm), we can speculate that this might be a contribution of a more significant number of disordered regions in CnDGRASP secondary structure (Figure 1C). We reconstructed the CD spectra of both human DGRASPs based on their PDB files by using PDB2CD and the reference database for soluble proteins SP175 (Figure S1) [37]. The reconstructed CD spectrum of DGRASP55 agrees very well with the experimental one, suggesting that the overall structure is not affected by the crystallographic conditions and packing (Figure S1A). The same is not observed for the DGRASP65, where the reconstructed CD spectra are very different from the experimental one (Figure S1B), suggesting that the structure in solution might differ in some degree from the crystallographic ones, even considering the inherent limitations of this reconstruction technique. These data suggest that all GRASP domains conserve the overall pattern of DGRASP55 crystal structure with CnDGRASP having more disordered regions.

**Figure 1:**
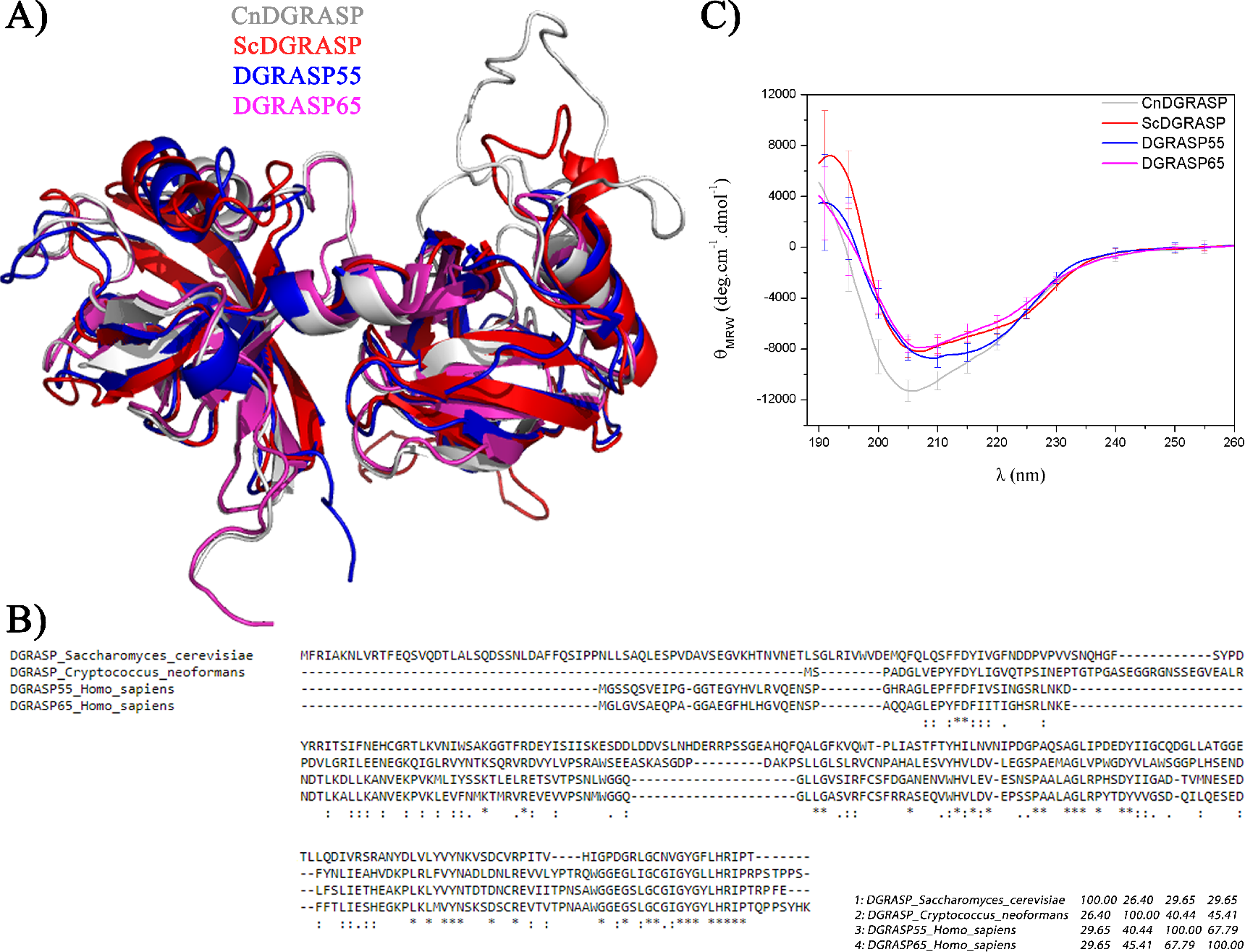
Structural analyses of different DGRASPs. A) Molecular modelling structures of CnDGRASP and ScDGRASP constructed using Swiss-Model, superimposed with the crystal structures of human DGRASP55 (PDB ID 3RLE) and rat DGRASP65 (PDB ID 4KFV). B) Sequence alignment of human DGRASPs, CnDGRASP and ScDGRASP using Clustal Omega. The Percent Identity Matrix is shown in the bottom right, where the numbers vary from 0 to 100 with 100 being the complete alignment of the sequences. C) Far-UV Circular dichroism spectra of the different GRASP domains.

The SRCD spectra deconvolution using Dichroweb confirmed our analyses by showing that the GRASP domains share about the same total secondary structure content (Table S1). The similarity observed in the CD spectra allows us to conclude that all GRASP domains used in this work share approximately the same secondary structure organization, although CnDGRASP seem to contain a greater contribution of disordered regions.

### 3.2 Looking for disordered regions within the domains

The SRCD deconvolution data indicated that all GRASP domains have considerable amounts of disordered regions (~40%). We previously reported the existence of intrinsically disordered regions (IDRs) in the full-length CnGRASP and ScGRASP, especially in the SPR domain [30]. Those conclusions were based on intrinsically disordered predictions and indirect experimental evidences, which impaired more definitive conclusions regarding IDRs in the GRASP domain only. IDRs are inherently sensitive to proteolysis and a proteolytic assay can be used to check for their indirect presence within the GRASP domains [50]. Here, trypsin was used in the limited proteolysis in a direct comparison with the stable and compact protein ovalbumin. As expected, Ovalbumin shows an incredible resistance to trypsin activity, a feature already observed previously (Figure 2) [30]. The fungi DGRASPs are very sensitive to this proteolysis assay (Figure 2) and it is possible to check that CnDGRASP is totally cleaved within 30 minutes of incubation, while ScDGRASP was promptly cleaved into two fragments that show some resistance to trypsin, (Figure 2). DGRASP55 shows the highest resistance to trypsin, with a total stability of at least 24 hours, similar to the ovalbumin control (Figure 2). Interestingly, it can be observed that a small fragment of this protein is cleaved after 30 minutes but the product is stable over time. We suggest that this might be a small disordered fragment from either the N-or the C-terminus, which can be cleaved without compromising the protein core. DGRASP65 still shows some resistance (compared to the fungi DGRASPs) but in a much less extent when compared with its paralogue (Figure 2). The protein core remains stable up to 3 hours only, indicating higher IDR content (Figure 2). The results suggest that both CnDGRASP and ScDGRASP contain higher IDR contents than human DGRASPs and that DGRASP65 seems to be somewhat similar to this subgroup than the well-behaved DGRASP55.

**Figure 2:**
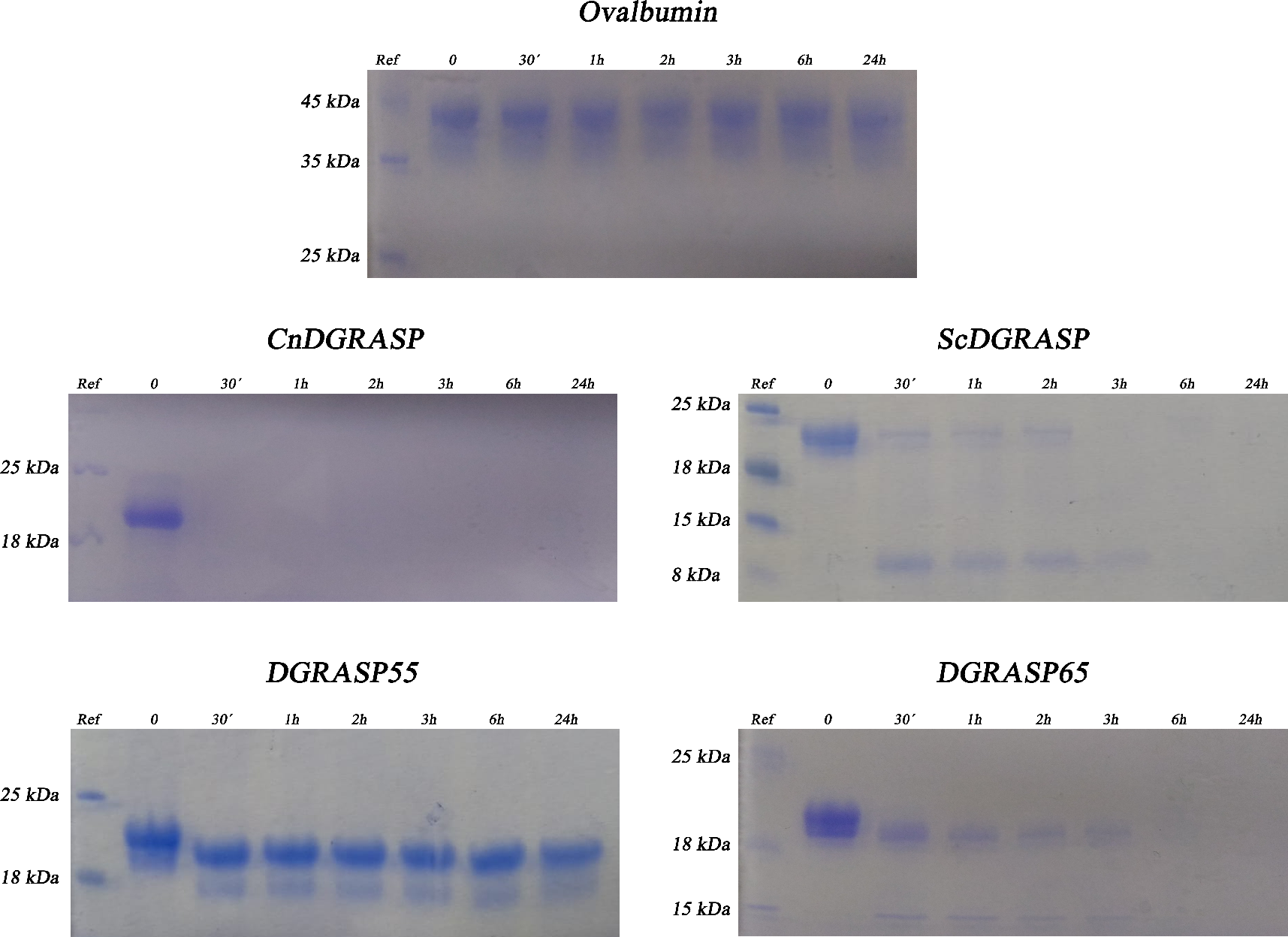
Proteolysis sensitivity monitored by SDS-PAGE. Different DGRASPs (concentration of 1 mg/mL) incubated with trypsin at a molar ratio of 1:100 for time intervals at 25 ℃. Ovalbumin, at the same protein:trypsin molar ratio and temperature, was used as a control of well-folded protein.

Another strategy to indirectly probe whether a protein presents an IDP-like behavior is by checking the effects of dehydration on its secondary structure, which can be monitored by CD or SRCD spectroscopies [50,51]. Unlike IDPs, globular, well-structured proteins exhibit only minor alterations by CD upon removal of bulk water during film formation [51]. We previously observed that the full-length CnGRASP shows significant structural changes upon dehydration, suggesting that the removal of water molecules can induce multiple disorder-to-order transitions [34]. However, no systematic study on the effects of dehydration over the GRASP domains has been performed. The CD spectra obtained from samples containing the GRASP domains under investigation before and after dehydration are shown in Figure 3. ScDGRASP and CnDGRASP show dramatic changes on the shape of their CD spectra when the bulk water is removed, where an increase of the *θ*_220 *nm*_ /*θ*_208 *nm*_ llill ratio and a significant shift of the peak at 190 nm to ~195 nm are observed (Figure 3). Both changes suggest an increase of the total regular/ordered secondary structure content [52]. Although the DGRASP65 shows higher exposure of their disordered regions when compared to DGRASP55, both structures are resistant to the dehydration effects, showing only minor changes after protein film formation (Figure 3). The data suggest that, even though all the DGRASPs chosen in this study present high disordered content (up to 40%), a difference in the packing/total tertiary interactions might expose (or not) these structures to the solution, leading to different accessibility to proteolytic activity and/or disordered-to-order transitions upon reducing the bulk water molecules content.

**Figure 3:**
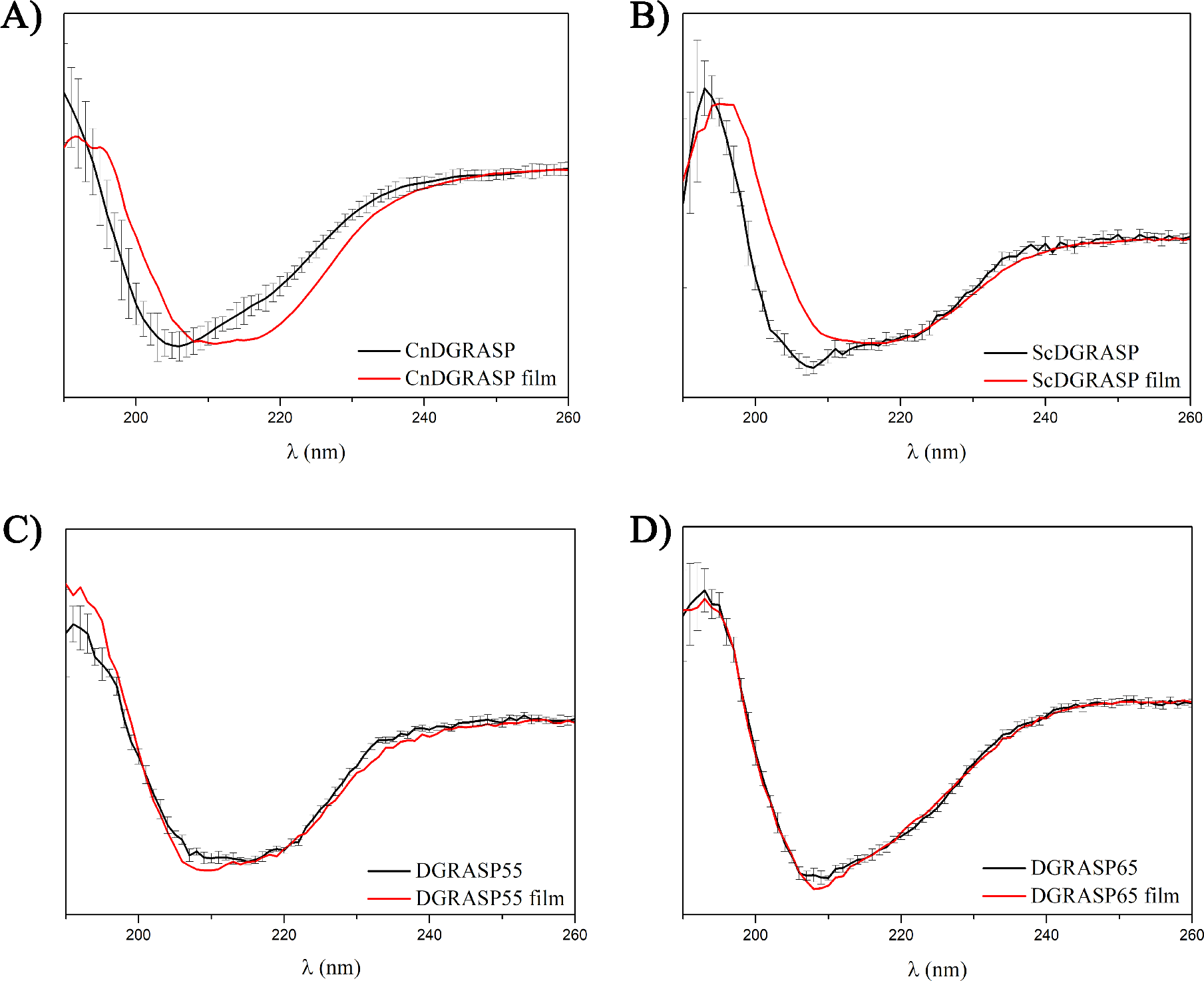
Secondary structure perturbation after dehydration monitored by SRCD and CD. In each panel, the red line is the spectrum from dried films of the proteins and the black, the control in solution. The error bars indicate one standard deviation in repeated measurements.

### 3.3 DGRASP55 is a more well-behaved protein

The simplest thermodynamical and kinetic models for a protein unfolding assume the protein can exist in two conformational states (native and unfolded), separated by a single activation free energy barrier, and inter-converting via a cooperative two-state transition [53]. Therefore, the degree of cooperativity can be used to check how well folded the protein is since this “well-behaving” is characterized by funnel-like energy landscapes that have a well-defined global energy minimum [53,54]. IDPs and “not so well-behaved” proteins lack this characteristic global minimum and the unfolding takes place in a less cooperative (or none) way due to multiple intermediate states [54,55]. Thus, one can look for IDP behavior by following the unfolding of a determined protein and checking the cooperativity of the transition, for instance using CD/SRCD experiments.

The chemical unfoldings of the GRASP domains are shown in Figure 4, where the ellipticities at characteristic wavelengths of the CD spectra are monitored as a function of urea concentration. The data suggest that, even though the fungi DGRASPs possess comparable amounts of ordered secondary structures compared to the human DGRASPs, their transition to the unfolded state takes place in a low cooperative way. ScDGRASP shows some degree of cooperativity but still very far from that observed in DGRASP55. It is possible to observe that DGRASP55 unfolding shows the characteristic sigmoidal transition, typical of well-behaved proteins [53] with a concentration of half transition centered around 4 M of urea. The data also indicate that DGRASP55 is a stable, well-folded, protein, which agrees with the fact that this protein is capable of forming well-diffracted crystals. However, DGRASP65 does not follow the same pattern (Figure 4). Although the crystal structure of apo human DGRASP65 is not yet available, there are structures solved for its apo and peptide-bound rat homologue [56,57], suggesting that we could expect a behavior similar to DGRASP55. However, this is not observed in Figure 4 and the transition to the unfolded state is linear, just like the one observed before for the full-length CnDGRASP [30]. The data indicate that the overall tertiary contacts of DGRASP65 share some of the structural features of the lower-eukaryotes DGRASPs, which were previously attributed to be collapsed IDPs/molten globule-like proteins [30,35].

**Figure 4:**
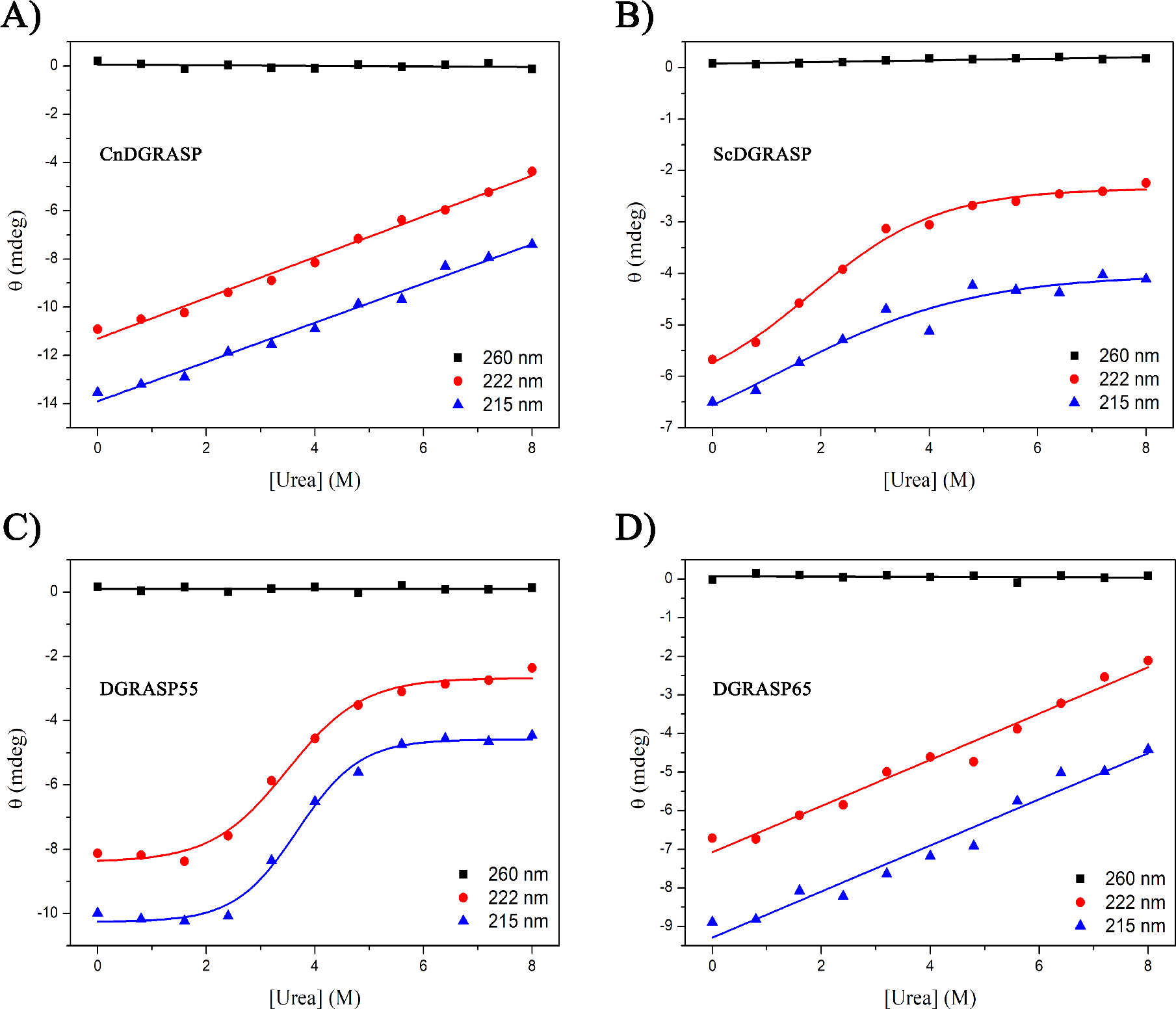
DGRASPs chemical denaturation using urea as a chaotropic agent. The graphics are showing the variation in ellipticity (mdeg) at different wavelengths (260, 222, 215 nm) during urea titration. A linear function was fitted to the data at 260 nm, representing the stability of the baseline, and to the 220/215 nm of CnDGRASP and DGRASP65. A sigmoidal function was fitted to the 222/215 nm of DGRASP55 and ScDGRASP.

### 3.4 DGRASPs have high IDR content but they are still collapsed proteins

Although rich in IDRs, all the GRASP domains used in this work have collapsed structures and this is easily observed by the maximum of tryptophan fluorescence emission, which are all blue shifted in some degree compared to the emission observed for the tryptophan in solution (Figure S2). Furthermore, the hydrophobic core is not easily accessible to the solvent, since the ANS permeation is not significantly high when compared to the unfolded-state in 8 M urea (Figure 5A). Native CnDGRASP is the only one showing high ANS accessibility (Figure 5A), a phenomenon already observed for the corresponding full-length construction [30]. As it has been shown previously, CnGRASP has a characteristic dynamic conformation in a μs-ms timescale leading to a hydrophobic interior with high water accessibility [30], and we show here that this behavior is also SPR-independent. The reason why only CnDGRASP shows this feature is not known but the higher IDR content could expose more hydrophobic regions, which might explain this behavior.

**Figure 5:**
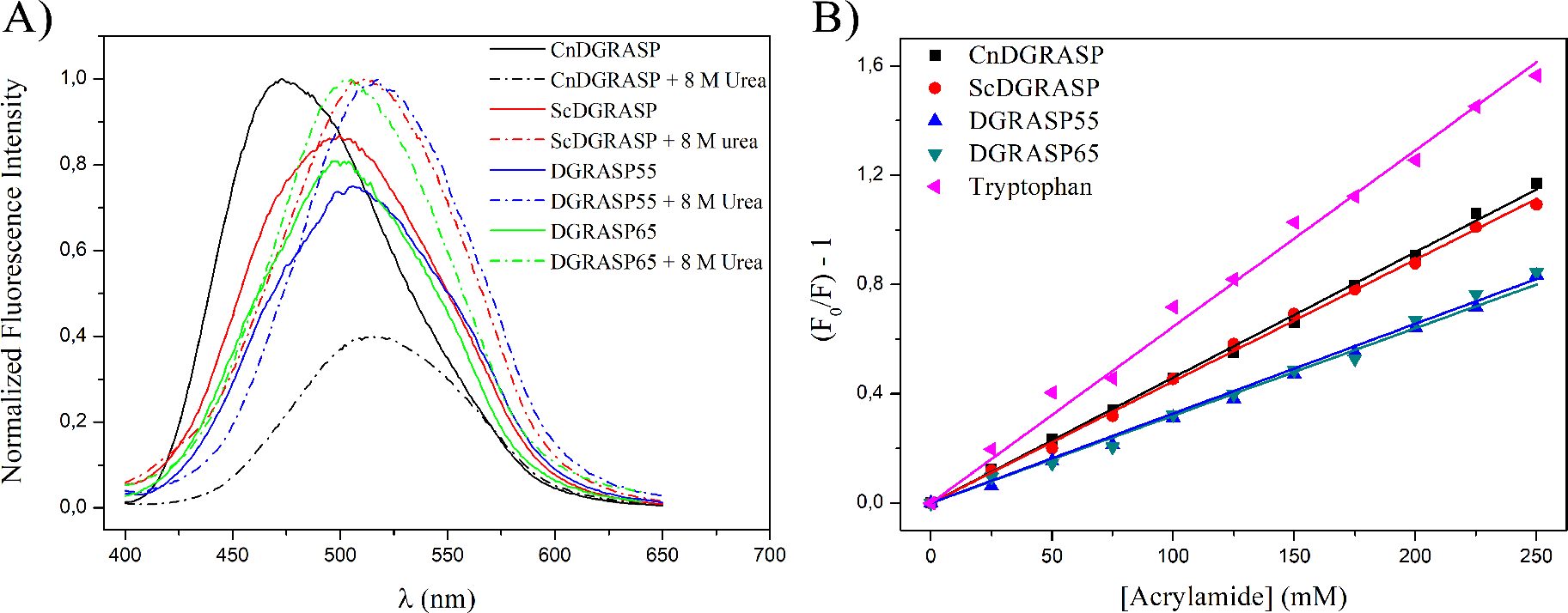
Protein compaction monitored indirectly using steady-state fluorescence. A) DGRASPs ANS accessibility. Each pair of spectra (with and without 8 M urea) was normalized dividing both dataset by the highest fluorescence intensity. This procedure indicates that only CnDGRASP has an accessible hydrophobic core in the native state. B) Fluorescence quenching using acrylamide. The linear Stern-Volmer model was fitted to each dataset and compared to the behavior of free tryptophan in solution.

CnDGRASP is the only DGRASP tested that has water-exposed hydrophobic sites, which do not include the regions where the Trp residues are located (Figure 5B). The tryptophan accessibility experiment data using the collisional quencher acrylamide agrees with the ANS experiment and show that all DGRASPs have a collapsed structure (different accessibility compared to the free tryptophan in solution). Trp residues present the same accessibilities to acrylamide between the two fungi DGRASPs and between the two human DGRASPs. The latter ones also show more collapsed and less accessible protein interior than the fungi ones (Figure 5B).

### 3.5 High-resolution data confirm that DGRASP65 and 55 are essentially different from each other

Although protein crystallography is not a suitable technique to tackle IDP structural determination, flexibility trends can still be inferred from the temperature factor (B-factor) of the crystal structures [50]. The B-factors of both DGRASP55 and DGRASP65 (PDB ID 3RLE and 4REY with the peptide-bounded omitted) are depicted on the respective structures in Figure 6A. The values observed for DGRASP55 are in the range 20-30 Å^2^, suggesting that this protein has a more well-defined structure. Those B-factor values are below the up B-factor maximum theoretical limit (around 31 Å^2^) acceptable for a structure solved at 1.6 Å resolution [58]. DGRASP65 B-factors show a different pattern, with the PDZ2 presenting a smaller B-factor range (similar to DGRASP55), but the PDZ1 being much more flexible, with several regions almost in the limit of total disorder (which is around 60 Å^2^), and much higher than the B-factor maximum expected for a structure solved at 1.9 Å resolution (around 40 Å^2^) [58]. Of course, this is not a suggestion that the crystal structures are right or wrong, but rather just an observation that DGRASP65 structure has a tendency of being more flexible than DGRASP55.

**Figure 6:**
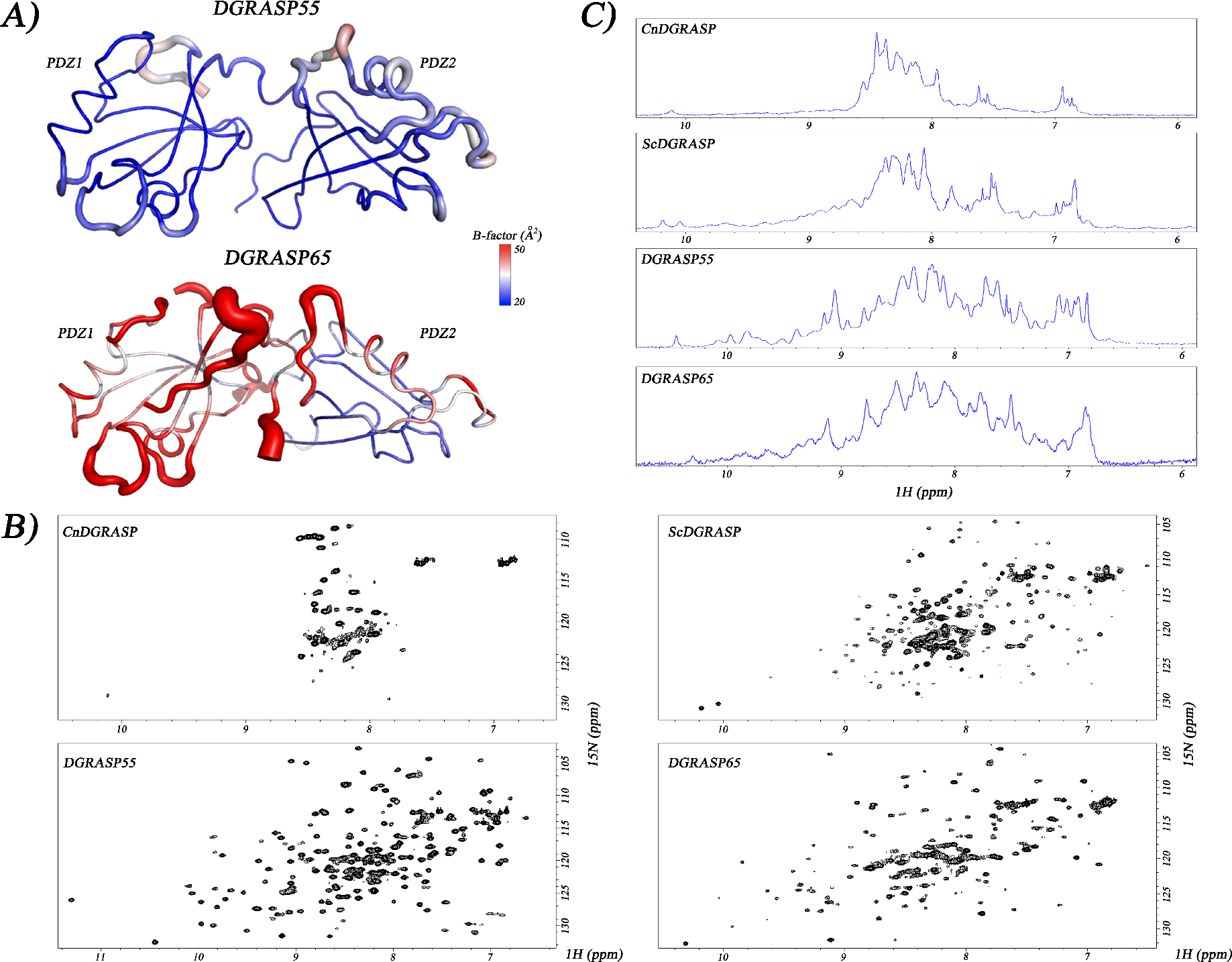
High-resolution analyses of DGRASP flexibility and disorder. A) Plot of the crystallographic structure of human DGRASP55 (PDB ID 3RLE) and human DGRASP65 with a peptide bounded which was omitted (PDB ID 4REY), in a color gradient according to their respective B-factor values. B) 1H-15N HSQC NMR spectra of DGRASPs. C) DGRASPS 1H NMR spectra of 1H-15N coupled protons.

Our low-resolution structural characterization presented above indicated that differences between DGRASPs are likely due to differences in IDR content. To further explore this issue, we used NMR to characterize the overall structure of these domains in solution. The ^1^H-^15^N HSQC spectrum of DGRASP55 shows that this domain is indeed well-folded and has low IDR content, since most of the amide resonances are widely spread from ca. 7 to 10 ppm (Figure 6B). DGRASP65 is again different with its ^1^H-^15^N HSQC spectrum showing several superpositions of amide resonances in the region around 8 ppm and narrower dispersion of the Asn/Gln side chains than that seen for DGRASP55 (Figure 6B). Nonetheless, DGRASP65 still shows good signal dispersion in its ^1^H-^15^N HSQC spectrum, suggesting that, although this protein is enriched in IDRs, it still possesses good amount of well-ordered structures.

The ScDGRASP ^1^H-^15^N HSQC spectrum is quite similar to the DGRASP65 but the signal crowding around 8 ppm is higher for the former, suggesting that ScGRASP is formed by an even higher number of IDRs (Figure 6B). This can be seen clearer in the proton spectrum where the majority of the resonance lines seems to be located (and overlapped) in this very narrow region around 8 ppm, especially when compared with the DGRASP65 or DGRASP55 proton spectra (Figure 6C). Again, CnDGRASP is the most different since the broadening of most of the resonance lines leads them to be out of the sensitivity limit and the few visible peaks in the HSQC NMR spectrum are located around 8 ppm (Figure 6B). This narrow distribution of resonance lines is more evident when looking to the proton resonance spectra in this range of the amide resonances (Figure 6C), where both fungi DGRASPs show similar spreading. The amide resonance line broadening follows the same pattern observed previously for the full-length CnGRASP, which was shown to be a molten-globule like protein with a conformational change-dynamic in a μs-ms timescale [34]. These data suggest that the CnDGRASP molten globule-like behavior is SPR independent.

### 3.6 DGRASPs show different behavior upon pH variation

IDPs are naturally disordered due to a greater/lower content of charged/hydrophobic residues, respectively [54,59]. It has been shown before that GRASPs do not have a combination of residues typical of fully disordered proteins [30], which agrees with our observations that they do possess ordered structures and are collapsed proteins. However, the presence of IDRs still leads to some unusual behavior and this might be due to uncompensated charges along those regions. In order to test this hypothesis, we evaluated how the DGRASPs respond to pH-induced charge alterations in their structures by measuring their CD spectra at different pH values (Figure 7). The secondary structure of CnDGRASP and DGRASP65 seems to be more resistant to pH variation, with only minor changes (within the error bars of the spectra) at higher pHs. CnDGRASP CD at pH 5 seems to be the only one with a change in shape, especially close to 205 nm. This was also observed previously for the full-length CnGRASP and was associated to the fact that this is closer to the theoretical pI of this protein, which is estimated to be 4.9. DGRASP55, however, shows significant changes at low pH (3 and 4). Because the pI of this protein is also around 5, these structural changes are just associated to a regular unfold. ScDGRASP is the one which has the most different and unusual pattern. At pH 3-4, this protein has a huge conformational change to a β-sheet rich structure, previously observed for the full-length ScGRASP and associated with the formation of a fibril-like supramolecular structure [35]. This data suggest that fibril formation might also take place with the isolated ScDGRASP at lower pHs, although this behavior was not observed for any other GRASP so far and it seems to be unique to yeasts.

**Figure 7:**
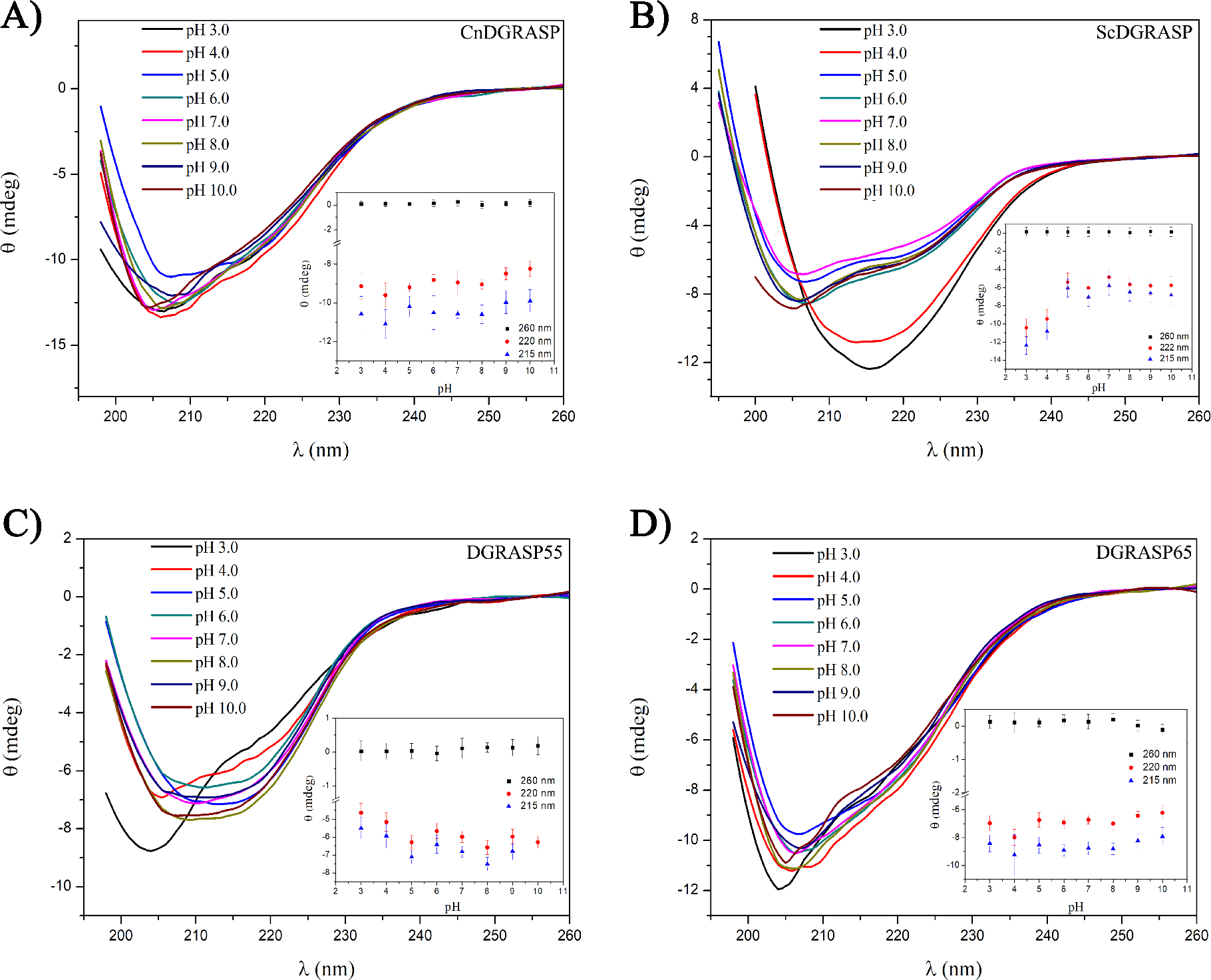
CD spectra of the GRASP domains at different pH values. The insets present the variation of the ellipticity at 260, 222, and 215 nm for each pH.

### 3.7 DGRASP65 is sequentially more similar to the fungi DGRASPs than DGRASP55

Eukaryotic diversity can be classified into six major divisions, or supergroups, based on phylogenomic analyses and named Opisthokonta, Amoebozoa, Excavata, Archaeplastida, Rhizaria and Chromalveolata [60]. The familiar model organisms of yeast and mammals are found within the Opisthokonta [60]. The Golgi apparatus is an ancient and ubiquitous organelle in eukaryotic cells [60]. Interestingly, it has been suggested before, based on a Golgi evolution phylogenetic analyses, that a GRASP65-homologue is likely to have been present in the last eukaryotic common ancestor since a GRASP65-homologue was observed in all 6 eukaryotic supergroups, although not in all the genomes tested [61]. We decided to analyze the sequence similarity between the DGRASPs (SPR-free sequence) in the human and the fungi, inside the Opisthokonta supergroup context, to look for correlations between the experimental data and protein sequence evolution. Our experimental data has shown that DGRASP65 shares characteristic from both DGRASP55 and fungi DGRASPs. The phylogenetic analyses support these observations (Figure 8). The evolutionary history suggested by the phylogenetic tree supports the expected clustering between species and DGRASPs, especially the groups of DGRASP55 and DGRASP65 (Figure 8A). Interestingly, the two outliers of the fungi DGRASPs tested are the ones used in this work, where CnDGRASP seems to be evolutionary closer to the mammalians than the other fungi DGRASPs and the ScDGRASP is the most different DGRASP (Figure 8A). To get information of which subgroup of mammalian DGRASPs is closer related to the fungi, we rearranged the bootstrap tree and forced a situation where the roots are in both human DGRASPs (Figure 8B and C). This was done only to force a situation where the opposite mammalian subgroup would collapse with the fungi. When the root is on the human DGRASP55, the subgroup of DGRASP65 and the fungi DGRASPs are collapsed in a node with a bootstrap of 95, given a high statistical significance to this arrange (Figure 8B). However, when the root is in the human DGRASP65, the two remaining subgroups collapse in a node with a bootstrap of only 55, so it is not possible to be certain about this organization (Figure 8C). These analyses suggest that the DGRASP65 subgroup seems to be closer than the DGRASP55 to the fungi DGRASPs in evolutionary terms. It is reasonable to speculate, from this sequence conservation, that GRASP65 might have appeared earlier in the evolution and, probably latter in mammalians, there was a GRASP65 duplication within the genome, giving rise to its paralogue GRASP55. It seems that the single fungi DGRASPs and DGRASP65 have evolved to keep a higher number of IDRs and a less packed structure, while DGRASP55 is more well-defined and well-behaved. The reasons why they are like this are still completely obscure.

**Figure 8:**
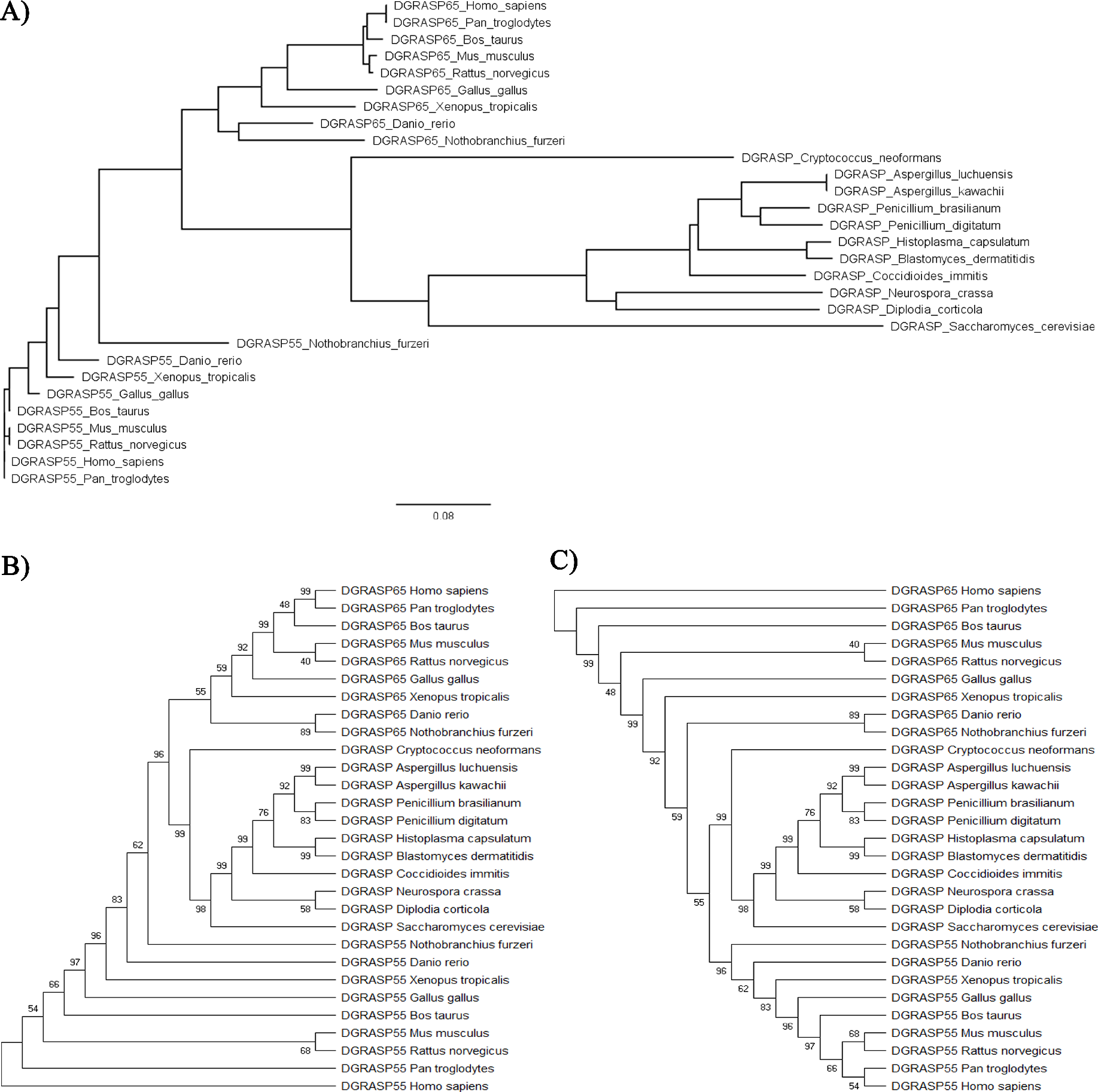
The evolutionary history inferred by using the Maximum Likelihood method and JTT matrix-based model estimated using Mega X. A) Phylogenetic tree with distance-scaled branch lengths constructed using FigTree v1.4.4. The MEGA X bootstrap consensus trees rooted in the B) DGRASP55 and C) DGRASP65 are also presented. The sequences used in this analysis are presented in the supplementary material.

## 4. Conclusions

GRASPs are fundamental part of the Golgi matrix and evolved to be not only a structural factor, but also a dynamic protein in other biological functions, especially UPS [9,19]. The GRASP domain is formed by two PDZ domains and this type of globule has been long described in the literature [14,31]. However, eukaryotic GRASPs are formed by PDZ domains presenting a unique fold [46]. We have previously observed unusual biophysical features of members of the GRASP family and it seems that IDRs play an important role in the structural plasticity of those proteins [30,34,35]. Nevertheless, how general is this behavior within the GRASP family and the role of the GRASP domain in these unusual biophysics observed for the full-length GRASP, are still a matter of debate. Here, we focused on four different GRASP domains: the two paralogues in humans (DGRASP55 and DGRASP65), the *S. cerevisiae* ScDGRASP (since most of the studies on UPS comes using this yeast as a model system) and the human-pathogen *C. neoformans* CnDGRASP.

Our results suggest that all those DGRASPs share the same fold previously observed in the crystal structure of human DGRASP55, with CnDGRASP showing a significant higher number of disordered regions (Figure 1). Our data also suggest that both fungi DGRASPs, which are also the solely GRASP genes found in the genome of those organisms, are enriched in IDRs and have low tertiary contacts (Figure 1, 2, 3, 4 and 6), making their transition to the unfolded state much less cooperative than those observed for general “well-behaved” proteins. This higher IDR content makes these proteins not only more sensitive to proteolysis (Figure 2), but also more prone to disorder-to-order transitions (Figure 3). They also show some interesting and unique characteristics, such as CnDGRASP having a higher ANS accessibility (Figure 5), which suggests an greater exposure of its hydrophobic core, and ScDGRASP being capable of forming fiber-like structures under some conditions (like pH, in this work, and temperature/dielectric constant variation in [35]).

On the other hand, the human DGRASP55 is the opposite of fungi DGRASPs and present itself as a more well-structured and well-behaved protein. DGRASP55 unfolding transition takes place in a high cooperative way, does not show significant disorder-to-order transitions upon dehydration and it is very resistant to proteolytic activity. Interestingly, DGRASP55 also shows high and impressive stability over time with no clear aggregation or unfold after a couple of weeks at 4c as monitored by size exclusion chromatography and CD spectroscopy (data not shown). It is worth noting how different DGRASP55 and the fungi DGRASPs are in biophysical means, even though they seem to share a similar fold. However, DGRASP65 seems to be placed in an intermediate state between DGRASP55 and the fungi DGRASPs and present higher content of IDRs compared to DGRASP55, but still less than the fungi DGRASPs, which leads to an intermediate proteolysis sensitivity and no effects upon dehydration. Its unfolding also presents a very low cooperative transition, similar to what is obtained for the CnDGRASP.

NMR was used to give higher resolution information on these IDRS and the data thus obtained agree with all the aforementioned results. The alignment of the primary sequences shows that DGRASP65 seems to be closer to the fungi DGRASPs in that respect. We hypothesize that the last common ancestor of fungi and humans, probably where the whole Opisthokonta supergroup diverged from the others, presented a GRASP homologue that shared the characteristics of ScGRASP, CnGRASP and GRASP65. Later in evolution, a gene duplication led to the paralogue GRASP55 appearance and this protein evolved to be a more well-structured protein for some still unknown reason. Probably this was the same evolutionary pressure that acted on the *Plasmodium falciparum* gene to evolve for two GRASP isoforms coming from an alternate splicing [17], one similar to GRASP55/65 and the other to the fungi ones.

This higher similarity between GRASP65 and the fungi GRASPs is reasonable if one thinks within the Golgi structure context, since most of the fungi have unstacked (or only partially stacked) Golgi complex and, in the reference of an unstacked, free cisternae, facing the ER, this would be more similar to the *cis*-Golgi face in mammalian cells. Since GRASP65 is present mainly in the *cis*-Golgi structure, while GRASP55 populates the *medial* and *trans* faces, it makes some sense to observe more similarity between GRASP65 and the fungi ones.

## Acknowledgments

The authors thank the Brazilian agencies Conselho Nacional de Desenvolvimento Científico e Tecnológico (CNPq) and Fundação de Amparo à Pesquisa do Estado de São Paulo (FAPESP) for the financial support through Grants No. 2015/50366-7, 2012/20367-3, and 308380/2013-4. LFSM and NAF acknowledge FAPESP for the post-doctoral (Grant No. 2017/24669-8) and PhD fellowship grants (Grant No. 2016/23863-2), respectively. JLSL is grateful for the financial support 303513/2016-0 from the CNPq-Brazil. The authors also thank the Multiuser Center for Biomolecular Innovation (EMU-Fapesp Grant No. 2009/53989-4) for the access to the NMR machine, and to ASTRID2 synchrotron at University of Aarhus, Denmark, for the access to the AU-CD beamline (beamtime grants to JLSL).

## Author contributions statement

LFSM performed most of the experimental procedures and the analyses. NAF, MCLCF, CGO, JLSL and FAM were involved in part of the experimental procedures and/or analyses. AJCF conceived and coordinated the project. All authors contributed to the writing and reviewing of the results. All authors also approved the final version of the manuscript.

