## Supplemental figures for "The GRASP domain in Golgi Reassembly and Stacking Proteins: differences and similarities between lower and higher Eukaryotes"

**Supplementary material**

**Understanding the structural conservation and features of the GRASP domain in “Golgi Reassembly and Stacking Proteins” from different Eukarya complexity**

Luís F.S. Mendes^1^, Natalia A. Fontana^1^, Carolina G. Oliveira^1^, Marjorie C.L.C Freire^2^, José L.S. Lopes^3^, Fernando A. Melo^4^, Antonio J. Costa-Filho^1^

1- Departamento de Física, Faculdade de Filosofia Ciências e Letras de Ribeirão Preto, Universidade de São Paulo, Ribeirão Preto, SP, Brazil

2- Centro de Pesquisas Aggeu Magalhães - Fundação Oswaldo Cruz (FIOCRUZ- PE)

3- Departamento de Física Aplicada, Instituto de Física, Universidade de São Paulo, SP, Brazil

4- Departamento de Física, Centro Multiusuário de Inovação Biomolecular, Universidade Estadual Paulista Júlio Mesquita, SP, Brazil

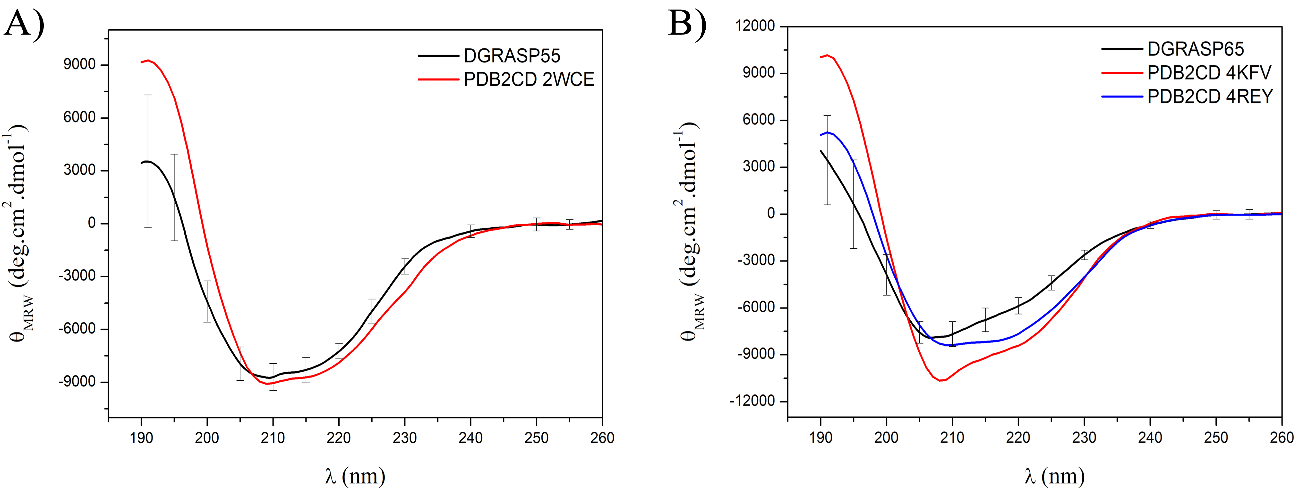

Figure S1: Circular Dichroism spectra reconstruction based on the PDB structure files were performed using PDB2CD. A) DGRASP55 experimental CD spectrum (black) with the theoretical CD based on the PDB file 2WCE (red). B) The same plot for DGRASP65 (black) but using the PDB files 4KFV (Rattus norvegicus homologue - red) and 4REY (crystal structure of the GRASP65-GM130 C-terminal peptide complex - blue). The GM130 C-terminal peptide was removed from the PDB file prior the CD spectrum reconstruction.

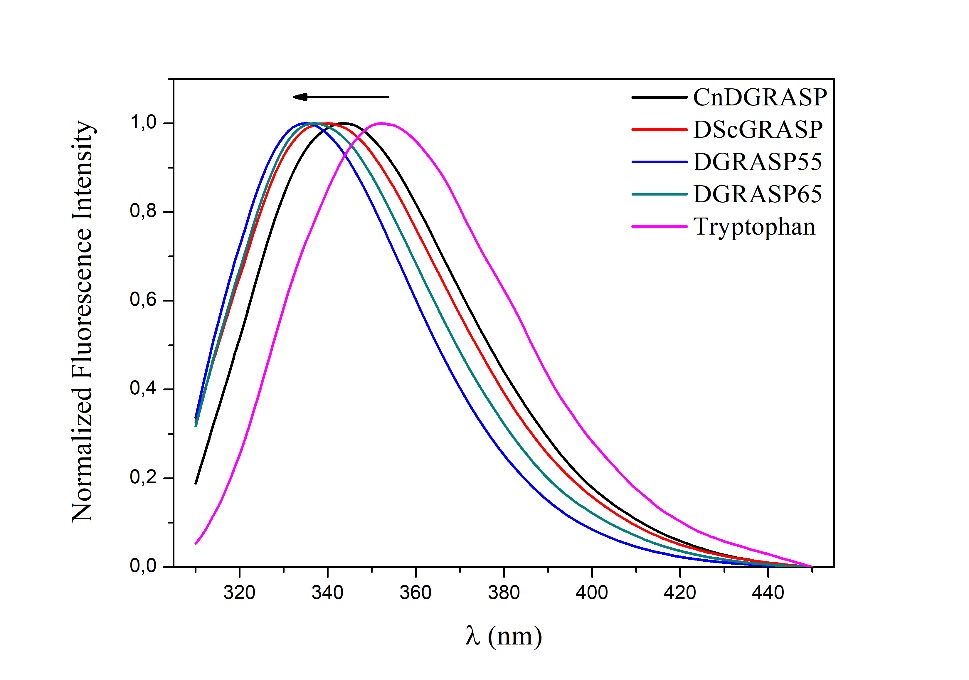

Figure S2: Steady-state fluorescence spectra of DGRASPs. The tryptophan selectivity excitation wavelength of 295 nm was used and the spectra acquired at 25$℃$. A L-Tryptophan solution (10 µM) was used as a control, and prepared using the DGRASPs buffer solution.

Table S1: CD spectra deconvolution using Dichroweb.

|  | **α-helix** | **β-sheet** | **Loop** | **Disordered** | **Total** | **NRMSD** |
| --- | --- | --- | --- | --- | --- | --- |
| ***CnDGRASP*** |  |  |  |  |  |  |
| - CDSSTR | 0.170 | 0.290 | 0.140 | 0.400 | 1.000 | 0.007 |
| - Selcom3 | 0.167 | 0.296 | 0.133 | 0.407 | 1.004 | 0.339 |
| - Contin-LL | 0.182 | 0.287 | 0.130 | 0.408 | 1 | 0.036 |
| Average | **17%** | **29%** | **13%** | **41%** |  |  |
| ***ScDGRASP*** |  |  |  |  |  |  |
| - CDSSTR | 0.170 | 0.330 | 0.130 | 0.390 | 1.000 | 0.028 |
| - Selcom3 | 0.169 | 0.297 | 0.121 | 0.353 | 0.940 | 0.259 |
| - Contin-LL | 0.182 | 0.312 | 0.128 | 0.378 | 1.000 | 0.129 |
| Average | **17%** | **31%** | **13%** | **37%** |  |  |
| ***DGRASP55*** |  |  |  |  |  |  |
| - CDSSTR | 0.160 | 0.310 | 0.130 | 0.390 | 0.99 | 0.023 |
| - Selcom3 | 0.158 | 0.327 | 0.127 | 0.374 | 0.986 | 0.203 |
| - Contin-LL | 0.164 | 0.324 | 0.127 | 0.385 | 1 | 0.158 |
| Average | **16%** | **32%** | **13%** | **38%** |  |  |
| ***DGRASP65*** |  |  |  |  |  |  |
| - CDSSTR | 0.100 | 0.340 | 0.130 | 0.430 | 1.000 | 0.020 |
| - Selcom3 | 0.138 | 0.309 | 0.133 | 0.389 | 0.969 | 0.130 |
| - Contin-LL | 0.175 | 0.287 | 0.130 | 0.408 | 1 | 0.036 |
| Average | **14%** | **31%** | **13%** | **41%** |  |  |

List of DGRASPs used in this work presented in a fasta format:

>DGRASP55_Homo_sapiens

MGSSQSVEIPGGGTEGYHVLRVQENSPGHRAGLEPFFDFIVSINGSRLNKDNDTLKDLLKANVEKPVKMLIYSSKTLELRETSVTPSNLWGGQGLLGVSIRFCSFDGANENVWHVLEVESNSPAALAGLRPHSDYIIGADTVMNESEDLFSLIETHEAKPLKLYVYNTDTDNCREVIITPNSAWGGEGSLGCGIGYGYLHRIPTRPFE

>DGRASP65_Homo_sapiens

MGLGVSAEQPAGGAEGFHLHGVQENSPAQQAGLEPYFDFIITIGHSRLNKENDTLKALLKANVEKPVKLEVFNMKTMRVREVEVVPSNMWGGQGLLGASVRFCSFRRASEQVWHVLDVEPSSPAALAGLRPYTDYVVGSDQILQESEDFFTLIESHEGKPLKLMVYNSKSDSCREVTVTPNAAWGGEGSLGCGIGYGYLHRIPTQPPSYHK

>DGRASP65_Mus_musculus

MGLGASSEQPAGGEGFHLHGVQENSPAQQAGLEPYFDFIITIGHSRLNKENDTLKALLKANVEKPVKLEVFNMKTMKVREVEVVPSNMWGGQGLLGASVRFCSFRRASEHVWHVLDVEPSSPAALAGLCPYTDYIVGSDQILQESEDFFTLIESHEGKPLKLMVYNSESDSCREVTVTPNAAWGGEGSLGCGIGYGYLHRIP

>DGRASP55_Mus_musculus

MGSSQSVEIPGGGTEGYHVLRVQENSPGHRAGLEPFFDFIVSINGSRLNKDNDTLKDLLKANVEKPVKMLIYSSKTLELREASVTPSNLWGGQGLLGVSIRFCSFDGANENVWHVLEVESNSPAALAGLRPHSDYIIGADTVMNESEDLFSLIETHEAKPLKLYVYNTDTDNCREVIITPNSAWGGEGSLGCGIGYGYLHRIP

>DGRASP55_Bos_taurus

MGSSQSVEIPGGGTEGYHVLRVQENSPGHRAGLEPFFDFIVSINGSRLNKDNDTLKDLLKANVEKPVKMLIYSSKTLELRETSVTPSNMWGGQGLLGVSIRFCSFDGANENVWHVLEVESNSPAALAGLRPHSDYIIGADTVMNESEDLFSLIETHEAKPLKLYVYNTDTDNCREVIITPNSAWGGEGSLGCGIGYGYLHRIP

>DGRASP65_Bos_taurus

MGLGASAEQPAGGAEGFHLHGVQENSPAQQAGLEPYFDFIITIGHSRLNKENDTLKALLKANVEKPVKLEVFNMKTMKVREVEVVPSNMWGGQGLLGASVRFCSFRRASEHVWHVLDVEPSSPAALAGLRPYTDYVVGSDQILQESEDFFSLIESHEGKALKLMVYNSESDSCREVTVTPNAAWGGEGSLGCGIGYGYLHRIP

>DGRASP55_Rattus_norvegicus

MGSSQSVEIPGGGTEGYHVLRVQENSPGHRAGLEPFFDFIVSINGSRLNKDNDTLKDLLKANVEKPVKMLIYSSKTLELREASVTPSNLWGGQGLLGVSIRFCSFDGANENVWHVLEVESNSPAALAGLRPHSDYIIGADTVMNESEDLFSLIETHEAKPLKLYVYNTDTDNCREVIITPNSAWGGEGSLGCGIGYGYLHRIP

>DGRASP65_Rattus_norvegicus

MGLGASSEQPAGGEGFHLHGVQENSPAQQAGLEPYFDFIITIGHSRLNKENDTLKALLKANVEKPVKLEVFNMKTMRVREVEVVPSNMWGGQGLLGASVRFCSFRRASEHVWHVLDVEPSSPAALAGLRPYTDYIVGSDQILQESEDFFTLIESHEGKPLKLMVYNSESDSCREVTVTPNAAWGGEGSLGCGIGYGYLHRIP

>DGRASP65_Gallus_gallus

MGLGSSSELPDGGAEGFHVHGVQENSPAQQGGLEPFFDFIIAIGHTRLNKENNMLKDLLKANAEKAVKLEVYNIKTMKIREVEVVPSNMWGGQGLLGASVRFCSFQGANEHVWHVLDVEPASPAAVGGLQPYTDYIVGSDQILQESEDFFSLIESHEGKPLKLMVYNTEADSIREVVVTPNGAWGGEGSLGCGIGYGYLHRIP

>DGRASP55_Gallus_gallus

MGASQSVEIPGGGTEGYHVLRVQENSPGHRAGLEPFFDFIVSINGSRLNKDNDTLKDLLKANVEKPVKMLVYSSKTLELRETSVTPSNMWGGQGLLGVSIRFCSFDGANENVWHVLEVEPNSPAALAGLRPHSDYIIGADTVMNETEDLFSLIETHEAKPLKLYVYNTDTDNCREVVITPNSAWGGEGSLGCGIGYGYLHRIP

>DGRASP65_Danio_rerio

MGLTQSSGDGPEGGTEGYHVHGVQEDSPAERAGLEPFFDFIISIGHNRLNQENDMLKDLLKANVEKPVKMEVYSTKTMRMRELEVVPSNMWGGQGLLGASVRFCSFQGANENVWHVLDVESNSPAALAGLQEHSDFIVGADQVLQDSEDFFSLIEAHEGKPLKLLIYNTETDKCREVVVTPNGAWGGEGSLGCGIGYGYLHRIP

>DGRASP55_Danio_rerio

MGGSQSVEIPGGGSEGYHVLRVQENSPGHRAGLEPFFDFIVSINNTRLNKDNDTLKDILKASVEKPVKMQVYSSKTLELREATVTPSNMWGGQGLLGVSIRFCSFEGANENVWHVLEVEPNSPAALAGLRPHTDYIIGADTVMNESEDLFSLIETHEGKGLKLYVYNTDTDNCREVVITPNSAWGGEGSLGCGIGYGYLHRIP

>DGRASP65_Pan_troglodytes

MGLGVSAEQPAGGAEGFHLHGVQENSPAQQAGLEPYFDFIITIGHSRLNKENDTLKALLKANVEKPVKLEVFNMKTMRVREVEVVPSNMWGGQGLLGASVRFCSFRRASEQVWHVLDVEPSSPAALAGLRPYTDYVVGSDQILQESEDFFTLIESHEGKPLKLMVYNSKSDSCREVTVTPNAAWGGEGSLGCGIGYGYLHRIP

>DGRASP55_Pan_troglodytes

MGSSQSVEIPGGGTEGYHVLRVQENSPGHRAGLEPFFDFIVSINGSRLNKDNDTLKDLLKANVEKPVKMLIYSSKTLELRETSVTPSNLWGGQGLLGVSIRFCSFDGANENVWHVLEVESNSPAALAGLRPHSDYIIGADTVMNESEDLFSLIETHEAKPLKLYVYNTDTDNCREVIITPNSAWGGEGSLGCGIGYGYLHRIP

>DGRASP55_Xenopus_tropicalis

MGGSNSVEIPGGGTEGYHVLRVQENSPGHRAGLEPFFDFIVSINGIRLNKDNDTLKDLLKANVEKPVKMVVYSSKTLELRETSVTPSNMWGGQGLLGVSIRFCSFEGANENVWHVLEVEPNSPAALAGLRAHSDYIIGADTVMNESEDLFSLIETHEGKPLKLYVYNTDTDNCREVVITPNTAWGGEGSLGCGIGYGYLHRIP

>DGRASP65_Xenopus_tropicalis

MGLSMSLSSEPLEGGTEGYHVHGIQENSPAQSAGLEPFFDFIIAIGHARLNKENSMFKDLLKANAEKPVKLEVYNTKSMKVREVEVTPSNMWGGQGLLGASVRFCSFQGANEHVWHVLDVEPNSPAALAGLQSHTDYIVGSDQILQESEDFYALVEAHEGKPLKLLVYNTETDSCREVFVTPNSAWGGEGSLGCGIGYGYLHRIP

>DGRASP65_Nothobranchius_furzeri

MGLSQSSEAAEGGTYGYHVHGVQPNSPAEKAGLQPFFDFILSLGNSRLNEENEQLKEVLKANVEKAIKMEVYSTKTTRVRELEVVPSNMWGGQGLLGASVRFCSYQGASENVWHVLDVETSSPAALAGLQSYSDYIVGADQVLQDSEDFFSLIEAHEGKPLKLLVYSTATDSCREVMVTPNGAWGGEGSLGCGIGYGYLHRIP

>DGRASP55_Nothobranchius_furzeri

MGGSQSVQIPGGGTEGYHVLRVQENSPGNRAGLEPFFDFIISVCDTRLKRDNDTLKELLKVNVERPVKMQLYNSKTQTVRETTVTPSNTWGGQGLLGVSIRFCSFEGANQNIWHVLEIEPNSPAALAGLRPFIDYIIGADIAMHESEDLFSLVEEHEGKELKLYVYGTDTDNCREVVITPNSDWGGEGSLGCGIGYGYLHRIP

>DGRASP_Cryptococcus_neoformans

MSPADGLVEPYFDYLIGVQTPSINEPTGTPGASEGGRGNSSEGVEALRPDVLGRILEENEGKQIGLRVYNTKSQRVRDVYLVPSRAWSEEASKASGDPDAKPSLLGLSLRVCNPAHALESVYHVLDVLEGSPAEMAGLVPWGDYVLAWSGGPLHSENDFYNLIEAHVDKPLRLFVYNADLDNLREVVLYPTRQWGGEGLIGCGIGYGLLHRIPRPSTPPS

>DGRASP_Saccharomyces_cerevisiae

MFRIAKNLVRTFEQSVQDTLALSQDSSNLDAFFQSIPPNLLSAQLESPVDAVSEGVKHTNVNETLSGLRIVWVDEMQFQLQSFFDYIVGFNDDPVPVVSNQHGFSYPDYRRITSIFNEHCGRTLKVNIWSAKGGTFRDEYISIISKESDDLDDVSLNHDERRPSSGEAHQFQALGFKVQWTPLIASTFTYHILNVNIPDGPAQSAGLIPDEDYIIGCQDGLLATGGETLLQDIVRSRANYDLVLYVYNKVSDCVRPITVHIGPDGRLGCNVGYGFLHRIPT

>DGRASP_Aspergillus_luchuensis

MFGALNRFIGLDAEPTRQARAQSTSDNSFGFQVLRNKDAELPLEPWFDFIVGINGRLIEDPDPNLFATEVRNCAGSSVTFEVWSAKGQKTHTVSIPVPASNPDLGLALQLAPLSSTQNIWHVLSIPSPLSPAYRAGLLPHSDYIIGTPSGTLRGESALGELVEDHLNRTLVLWVYNSEFDVVREVELVPTRGWGGEGALGAELGFGALHRLPVGL

>DGRASP_Neurospora_crassa

MTTMFNALNRFISRLDGDAATSKENQPGGFGFQVLRNTNPDLAIEPWFDFVVGINGRMIDDSDPRLFAQEVRNCAGGVVQLGLWSAKGQRTRALHIPVPADTASLGLTLQWTALSVVTNIWHVLDVPANSPADVAGLLPYSDYILGTPEGVLHGESGLSELVEDHIDRPLRLYIYNNEYNVTREVTIQPSRDWGGEGALGCVLGYGALHRLPAP

>DGRASP_Penicillium_brasilianum

MFGALNRFIGRLDGEPVQQPRNGPSDNAYGFQVLRNKDTELPLEPWFDFIVGINGHPIEDPDPNLFATEVRNCAGSSLTLEIWSAKGQRTHTVTVPISATTPSLGLALQLAPLSSTQNIWHVLGIPSPLSPAFRAGLLPHSDYIIGTPSGTLRGESALGELVEDHLDRTLVLWVYNSEFDVVREVELIPTRGWGGEGALGAELGYGALHRLPIGL

>DGRASP_Coccidioides_immitis RS

MFGALNRFIGRLDSDLPKQPTSTTGDNSYGFQILRNKDPDLPLEPWFDFIVGINGRLIDEPDPHLFATEVRNCAGTSVSLEIWSAKGQRTHNVVIPIPPEKPTLGLTLQLAPLSSTQHIWHILSIPSPLSPAYLAGLLPHSDYILGTPSGTLRGEAALGELVEDHLNRSLTLWVYNSEFDVVREVEIVPNRNWGGEGALGAVLGYGALHRLPVG

>DGRASP_Diplodia_corticola

MSMFGALNRFISRLDAEPQDRDQGTKGAYGFQILRNTNQELSIEPWFDFIIGINGRTIDDPDPTLFATEVRNCAGHTISLGVWSAKGQRIRELYVPIPAETPTLGISLQWSPLSSTEDVWHILDVQPNSPADLAGLLPYGDYVIGSPEGLVRGESGLGELVEDYINRPLRLFVYNHEYNVTRPITITPSRNWGGQGLLGCVLGFGALHRVPAP

>DGRASP_Histoplasma_capsulatum H88

MFSALNRFIARLDSEPGQQQHSHFRDNPHGFQVLRNKDPELLLEPWFDFVIGINGHLIDDPDPNLFATEIRNCAGSSVTFEVWSAKGQRTYTTTIAVPSENPTLGVTFQFSPLSATQHIWHITEIPSPLSPAYQAGLLPHSDYILGTPSGTLRGESALGELVEDHLNRSLVLWVYNSEFDVVREVELVPRRGWGGEGALGAVLGYGALHRLPFG

>DGRASP_Blastomyces_dermatitidis

MFGALNRFIARLDSEPGQQQHSHFRDNPHGFQVLRNKDPELPLEPWFDFVIGINGHLIDDPDPNLFATEIRNCAGSSVTFEIWSAKGQRTYTTTVPVPSEKPTLGVTFQFSPLSTTQHIWHITEIPSPLSPAYQAGLLPHSDYILGTPSGTLRGESALGELVEDHLNRSLVLWVYNREFDVVREVELVPRRGWGGEGALGAVLGYGALHRLPVGLG

>DGRASP_Penicillium_digitatum

MFGALNRFISRLDGEPGQQPRNGPSDNAFGFQVLRNKDPELPLEPWFDFIVGINGHTIEDPDPNLFATECRNCAGGSVTLEVWSAKGQRTHTVTHPIPPTNPSLGLALQLAPLSSTHNIWHVIAIPSPLSPAYRAGLLPYSDYIIGTPSGTLRGESALGELVEDHLNRTLVLWVYNSEFDVVREVELVPTRGWGGEGALGAELGYGALHRLPIG

>DGRASP_Aspergillus_kawachii

MFGALNRFIGLDAEPTRQARAQSTSDNSFGFQVLRNKDAELPLEPWFDFIVGINGRLIEDPDPNLFATEVRNCAGSSVTFEVWSAKGQKTHTVSIPVPASNPDLGLALQLAPLSSTQNIWHVLSIPSPLSPAYRAGLLPHSDYIIGTPSGTLRGESALGELVEDHLNRTLVLWVYNSEFDVVREVELVPTRGWGGEGALGAELGFGALHRLPVGLGE
